# AlphaFold2-Guided Functional Screens Reveal a Conserved Antioxidant Protein at ER Membranes

**DOI:** 10.1101/2024.06.19.599784

**Authors:** Zhijian Ji, Taruna Pandey, Henry de Belly, Jingxuan Yao, Bingying Wang, Orion D. Weiner, Yao Tang, Shouhong Guang, Shiya Xu, Zhiyong Lou, Thomas D. Goddard, Dengke K. Ma

## Abstract

Oxidative protein folding in the endoplasmic reticulum (ER) is essential for all eukaryotic cells yet generates hydrogen peroxide (H_2_O_2_), a reactive oxygen species (ROS). The ER-transmembrane protein that provides reducing equivalents to ER and guards the cytosol for antioxidant defense remains unidentified. Here we combine AlphaFold2-based and functional reporter screens in *C. elegans* to discover a previously uncharacterized and evolutionarily conserved protein ERGU-1 that fulfills these roles. Deleting *C. elegans* ERGU-1 causes excessive H_2_O_2_ and transcriptional gene up-regulation through SKN-1, homolog of mammalian antioxidant master regulator NRF2. ERGU-1 deficiency also impairs organismal reproduction and behavioral responses to H_2_O_2_. Both *C. elegans* and human ERGU-1 proteins localize to ER membranes and form network reticulum structures. Human and *Drosophila* homologs of ERGU-1 can rescue *C. elegans* mutant phenotypes, demonstrating evolutionarily ancient and conserved functions. In addition, purified ERGU-1 and human homolog TMEM161B exhibit redox-modulated oligomeric states. Together, our results reveal an ER-membrane-specific protein machinery for peroxide detoxification and suggest a previously unknown and conserved mechanisms for antioxidant defense in animal cells.

## Introduction

In eukaryotic cells, the endoplasmic reticulum (ER) plays a critical role in protein folding, a process essential for cell function but one that generates the reactive oxygen species hydrogen peroxide (H_2_O_2_)^1–3^. The ER-residing oxidase Ero1α stoichiometrically produces one molecule of H_2_O_2_ per disulfide bond introduced to nascent secreted or membrane proteins. Accordingly, the H_2_O_2_ concentration in the ER lumen is estimated to be approximately 700 nM, whereas cytosolic H_2_O_2_ concentration is estimated to be 2.2 nM at steady levels^4–6^. Low cytosolic H_2_O_2_ levels are maintained by various H_2_O_2_-buffering systems and enzymes, including H_2_O_2_-degrading catalases. Known families of catalase localize to peroxisomes, cytosol and mitochondria, but not ER^7–10^. While H_2_O_2_ can cross membranes by slow diffusion or aquaporin-facilitated transport^11^, it remains unknown how ER membranes are guarded against the exceptionally high levels of ER H_2_O_2_, whose derivatives (e.g. hydroxyl radicals via Fenton reactions) can chemically attack and damage membrane lipids, nucleic acids and proteins.

In bacteria, a family of transmembrane proteins (DsbD/ScsB)^12^ provide reducing equivalents for correct disulfide bond formation and peroxide reduction in the periplasm, which is considered the bacterial equivalent of eukaryotic ER. DsbD and ScsB carry multi-transmembrane segments and employ a series of cysteine pairs with differential redox potentials to relay electron transfer from cytosol to periplasm^13^. Mammalian counterparts of DsbD and ScsB are proposed to exist yet remain unidentified^14^. Our study addresses this gap in knowledge by identifying a previously uncharacterized and evolutionarily conserved *C. elegans* protein *Y87G2A.13* (named as ERGU-1, <u>ER gu</u>ardian of oxidative stress) as the long-sought ER membrane-localized antioxidant protein. ERGU-1 protects against elevated H_2_O_2_ and ensures optimal organismal functions in *C. elegans*. Furthermore, we reveal the structural and functional conservation of ERGU-1 homologs across various animal species, indicating an ancient and fundamental role for ERGU-1 in maintaining cellular redox homeostasis.

## Results

### Computational and genetic discovery of ERGU-1

We first performed BLASTP search to identify potential eukaryotic sequence homologs of DsbD and ScsB. Such amino acid sequence similarity-based queries yielded no apparent broadly conserved eukaryotic homologs, even after adjusting sensitivity and specificity of the search (Fig. S1). We reasoned that DsbD/ScsB and potential eukaryotic counterparts might not share primary protein sequence similarities owing to distant evolutionary divergence. Nevertheless, eukaryotic counterparts of DsbD/ScsB could exhibit structural and functional, rather than protein sequence features, common for DsbD/ScsB and known families of transmembrane oxidoreductases. These features include multi-transmembrane segments to facilitate intramembrane electron transfer, closely spaced cysteine clusters in different transmembrane segments within proximity (5-10 Angstroms), and broad evolutionary conservation in eukaryotic or multicellular organisms. Based on these features, we sought to leverage the availability of predicted protein structures by AlphaFold2 for nearly the entire *C. elegans* proteome^15^ and filter through DsbD/ScsB-like candidates for subsequent functional screens and validation in *C. elegans*.

To identify DsbD/ScsB-like candidates, we conducted a computational search of the *C. elegans* proteome based on the functional features common for DsbD/ScsB-like transmembrane oxidoreductases. Using the *C. elegans* UniProt reference proteome as a starting point (version 26, 19,827 proteins), we found all genes with at least 4 annotated transmembrane helices (3,177 proteins) and further filtered to those with at least two transmembrane helices, each containing at least two cysteines separated by at most 2 intervening residues (sequence patterns CC, CXC, or CXXC). For the 190 proteins meeting these criteria, we examined AlphaFold2 Database-predicted structures and selected those in which cysteine pairs in two helices were no more than 10 Angstroms apart (SG to SG atom distance), producing 53 candidate genes (Fig. 1A and Fig. S1). We implemented the computational screen using customized Python scripts (Supplementary Data file) in UCSF ChimeraX^16^.

**Fig. 1.**
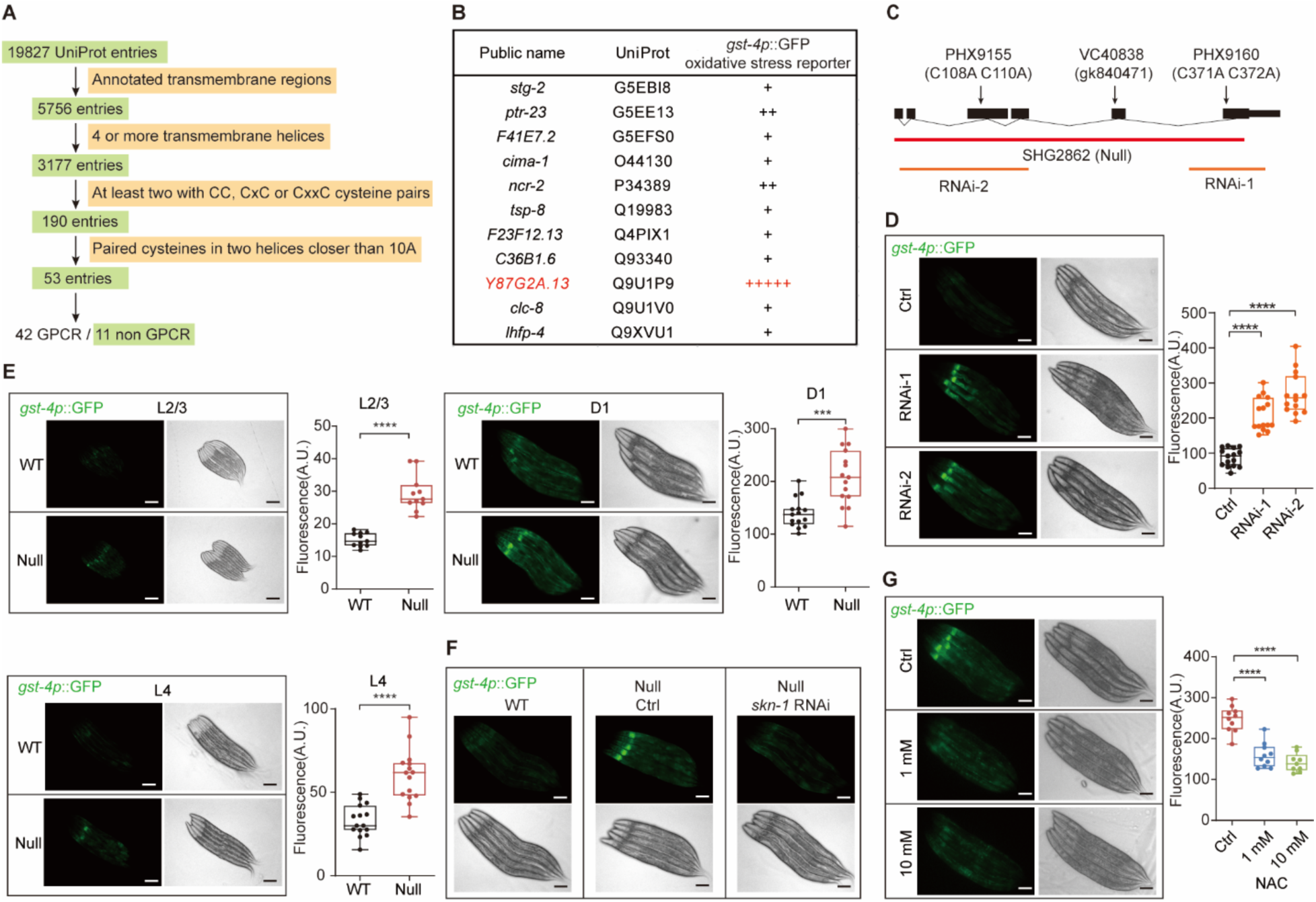
Computational and genetic identification of ERGU-1. (**A**) Schematic of AlphaFold2-assisted computational screens. (**B**) Phenotype-driven functional screens based on RNAi activation of *gst-4p::GFP* oxidative stress reporters. (**C**) Gene structure of *ergu-1* showing various alleles. (**D** and **E**) Representative epifluorescence images showing *gst-4p::GFP* activation in two independent *ergu-1* RNAi (D) and mutants (E) (n=15 for each group). (**F** and **G**) Representative epifluorescence images showing that *gst-4p::GFP* activation in *ergu-1* null mutants requires ROS and *skn-1* (F), which are suppressed by NAC dose-dependently (G) (n=10 for each group), with quantification of *gst-4p*::GFP fluorescence intensities under conditions indicated. *P* value was determined by an unpaired *t*-test, two-tailed (comparison between two groups) or one-way ANOVA (comparison between multiple groups). Scale bars, 100 μm.

Among the 53 candidates with AlphaFold2-predicted structures passing the above screening criteria, we focused on the non-GPCR category with 11 hits for functional validation in *C. elegans* (Fig. 1A, B). We performed RNA interference (RNAi) against genes encoding these 11 hits and tested if any can activate the transcriptional *gst-4p::GFP* reporter, which has been previously used to monitor excess oxidative stress and elevated levels of peroxide in *C. elegans*^17^. We found that RNAi against one gene *Y87G2A.13* strongly activated *gst-4p::GFP* under standard laboratory conditions without exogenous oxidative stress, while RNAi against other genes or empty vector control did not (Fig. 1B). Based on the T777T vector backbone with improved RNAi specificity and efficacy^18^, we designed two independent RNAi constructs targeting different coding regions of *Y87G2A.13* and obtained similar results (Fig. 1C, D). In addition, we used CRISPR-mediated genetic deletion to generate a null allele of *Y87G2A.13* and observed similar constitutive *gst-4p*::GFP activation in *Y87G2A.13* null mutants (Fig. 1C, E). We also observed similar *gst-4p*::GFP activation in *ergu-1(gk840471)* mutants with a protein-truncating mutation (Fig 1C, Fig. S2A, B). The transcription factor SKN-1 is the *C. elegans* ortholog of NRF2, master regulator of antioxidant responses in mammals, and mediates *gst-4p::GFP* activation upon a variety of oxidative stresses^19–21^. We found that RNAi against *skn-1* abolished the constitutive *gst-4p::GFP* activation in *Y87G2A.13* null and protein-truncating mutants (Fig. 1F, Fig. S2C). N-acetyl-cysteine (NAC), a commonly used precursor for glutathione-based antioxidants and ROS/H_2_O_2_ scavenger in *C. elegans*^22–24^, also strongly suppressed *gst-4p::GFP* activation in *Y87G2A.13* null mutants (Fig. 1G). Taken together, these results show that reduced expression by RNAi or genetic deficiency of *Y87G2A.13*, one of the computationally identified ERGU candidates, constitutively activates oxidative stress response in a manner that involves excess oxidants and activation of SKN-1. We hereafter refer to the Y87G2A.13 protein as ERGU-1 (<u>ER</u> <u>gu</u>ardian of redox defense), given additional lines of evidence below.

### Molecular and organismal roles of ERGU-1 in antioxidant defense in *C. elegans*

As intramembrane oxidoreductases, DsbD and ScsB transfer electrons via cysteine pairs as redox couples to reduce excess protein disulfide bonds and peroxides in bacterial periplasm^12^. We thus assessed similar roles of ERGU-1 in ER protein-folding stress and H_2_O_2_ reduction, using the *hsp-4p::GFP* transcriptional reporter and a specific fluorescent sensor roGFP2-Orp1 for H_2_O_2_, respectively^17,25^. We found that *ergu-1* genetic deletion or RNAi caused constitutive activation of *hsp-4p::GFP* in the absence of exogenous ER or protein folding stresses (Fig. 2A, Fig. S3A). Using *C. elegans* strains expressing roGFP2-Orp1 to specifically monitor cytosolic levels of H_2_O_2_ through ratiometric dual-color fluorescence (Fig. 2B), we found that *ergu-1* genetic deletion or RNAi caused markedly higher ratios of sensor oxidation versus reduction, indicating elevated H_2_O_2_ levels. An independent assay based on non-fluorescent compound dichlorofluorescein (DCFH) that reacts with H_2_O_2_ to produce fluorescent 2’,7’-dichlorofluorescein (DCF) showed similarly higher levels of peroxide stress in *ergu-1* mutants (Fig. 2C). Furthermore, we assayed effects of *ergu-1* null, protein-truncating mutants or RNAi on a comprehensive panel of stress-responding reporters, cell type-specific markers, and organelle membrane lipid sensors^26–28^, but did not observe apparent phenotypic consequences, except oxidative stress-related reporters *gst-4p*::GFP, *hsp-4p*::GFP, roGFP2-Orp1 and Grx1-roGFP2 (Fig. S2D, 3A-3M). These results suggest ERGU-1 normally defends against specific peroxide-related oxidative stress, consistent with its structural features and functional reporter regulation observed.

**Fig. 2.**
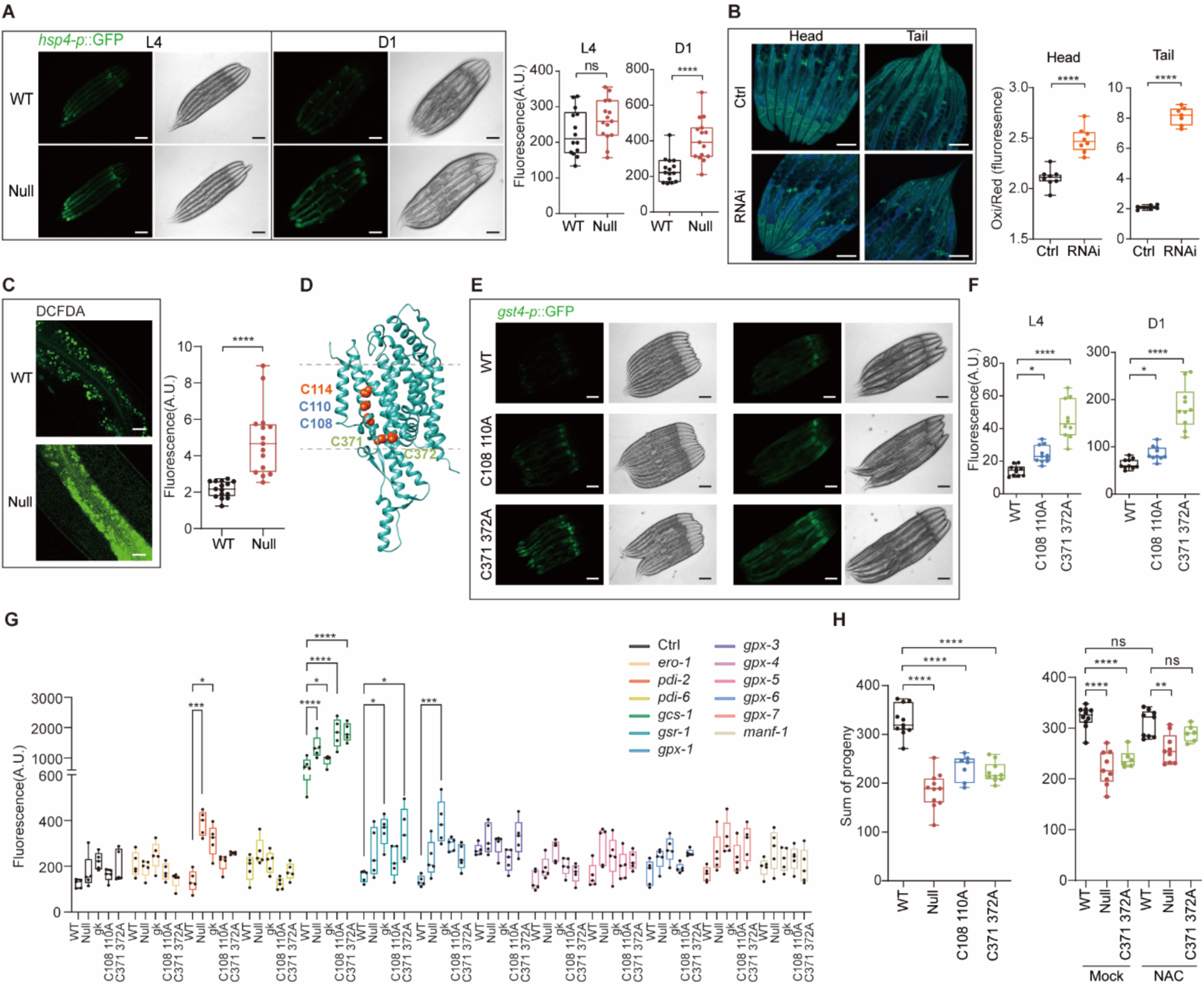
ERGU-1 defends against H_2_O_2_ and maintains organismal functions. (**A**) Representative epifluorescence images showing increased *hsp-4p::GFP* in *ergu-1* null mutants (n=15 for each group). (**B**) Representative confocal images showing increased oxidation/reduction ratio of Orp1-roGFP (H_2_O_2_ indicator) with *ergu-1* RNAi. (**C**) DCFDA staining showing increased DCF in null mutants (n=15 for each group). (**D**) AlphaFold2-predicted structure revealing cysteine residues that likely mediate electron transfer across membranes. (**E**) Representative epifluorescence images showing increased *gst-4p*::GFP in two different cysteine pair mutants. (**F**) Quantification of *gst-4p*::GFP fluorescence intensities in WT and two different cysteine pair mutants indicated (n=10 for each group). (**G**) Fluorescence intensity of *gst-4p*::GFP in wild type (WT) and *ergu-1* null mutants under control (Ctrl) and RNAi conditions targeting redox regulators, including *ero-1*, *pdi-2*, *pdi-6*, *gcs-1*, *gsr-1*, *gpx-1*, *gpx-3*, *gpx-4*, *gpx-5*, *gpx-6*, *gpx-7*, and *manf-1*. (**H**) Quantification of brood sizes in wild type, *ergu-1* null and cysteine mutants (left panel). Comparison of brood sizes between wild-type and mutant strains under mock and 10 mM NAC treatments (right panel). *P* value was determined by an unpaired *t*-test, two-tailed (comparison between two groups) or one-way ANOVA (comparison between multiple groups). [(A), (B) and (E)] Scale bars, 100 μm. (C) Scale bars, 20 μm.

The AlphaFold2-predicted structure of ERGU-1 reveals two clusters of cysteine pairs in its two close transmembrane segments (Fig. 1B, 3D). To examine the importance of such cysteine pairs for the antioxidant function of ERGU-1, we obtained CRISPR-mediated knock-in alleles converting cysteine to alanine at each site, and crossed such cysteine mutants separately with reporter strains of *gst-4p::GFP*. We found that these cysteine mutants caused elevated levels of *gst-4p::GFP* (Fig. 2E, F). In addition, we found that the *ergu-1* null mutant phenotypes were rescued by transgenic extrachromosomal arrays of wild-type but not cysteine mutants of *ergu-1* (Fig. S4A). These results suggest that the clusters of cysteine pairs of ERGU-1 functionally contribute to defending against oxidative stress, supporting their roles in relaying electron transfer across ER membranes to reduce peroxide stress and excess H_2_O_2_ from ER.

We next determined organismal roles of ERGU-1. Morphologically, the null and cysteine knock-in *ergu-1* mutants appear grossly normal. Previous studies have implicated roles of SKN-1 activation in several long-lived genetic mutants or by oxidative stress-induced hormesis in promoting longevity^19^. Unexpectedly, we found that the *ergu-1* null mutants, with control or *skn-1* RNAi, exhibited largely normal lifespans (Fig. S2H) under standard laboratory conditions at 20 °C fed with OP50 on FuDR-NGM plates. To test genetic interaction of *ergu-1* with other known genes implicated in ER-specific redox regulation, we examined effects of RNAi against genes in the *ero-1, gcs-1, gsr-1, gpx* and *pdi* families and found that *ergu-1* mutants showed increased oxidative stress responses compared with wild type upon RNAi treatment (Fig. 2G). Notably, assay quantification of reproductive capacity revealed a markedly reduced brood size of *ergu-1* null mutants. Consistently, the *ergu-1* cysteine mutants exhibited similar reduction in brood sizes (Fig. 2H). This reduced brood size phenotype can be partially alleviated upon 10 mM NAC antioxidant treatment (Fig. 2H). By contrast, treatment with exogenous oxidative stressor paraquat decreased brood size more severely in *ergu-1* mutants than wild type (Fig. S3I). In addition, we measured the locomotion speed change of young adult hermaphrodites upon exogenous H_2_O_2_ and found that *ergu-1* null mutants exhibited a markedly occluded locomotion-slowing response to H_2_O_2_, without apparently affecting neuronal development, cytoskeletal structure or baseline neuronal activity (Fig. S3J-S3N). These results indicate that ERGU-1 is critical for maintaining normal specific organismal functions in reproduction and behavioral responses to H_2_O_2_.

### Subcellular localization of *C. elegans* ERGU-1 at ER membrane

To elucidate the tissue distribution and subcellular localization of ERGU-1, we constructed both a transcriptional (*ergu-1p*::GFP, fusing the *ergu-1* promoter to the green fluorescent protein GFP) and a translational reporter (*ergu-1*p::*ergu-1*::GFP, fusing GFP to the C-terminus of ERGU-1 under the control of its native promoter) (Fig. 3A). We microinjected the constructs and integrated the transgenic extrachromosomal arrays to the genome at low copy number to ensure the faithful recapitulation of ERGU-1’s endogenous expression pattern. Both the transcriptional and translational reporters similarly revealed *ergu-1* expression in major metabolic tissues, including the intestine, body wall muscles, and the spermatheca (Fig. 3B, C). As expected, RNAi against *ergu-1* abolished GFP signals from the *ergu-1*p::*ergu-1*::GFP transgene (Fig. S3B). Notably, the tissues expressing *ergu-1* are also known to exhibit high metabolic activities, generating elevated levels of H_2_O_2_ during protein folding, as indicated by roGFP2-Orp1 (Fig. 2B).

**Fig. 3.**
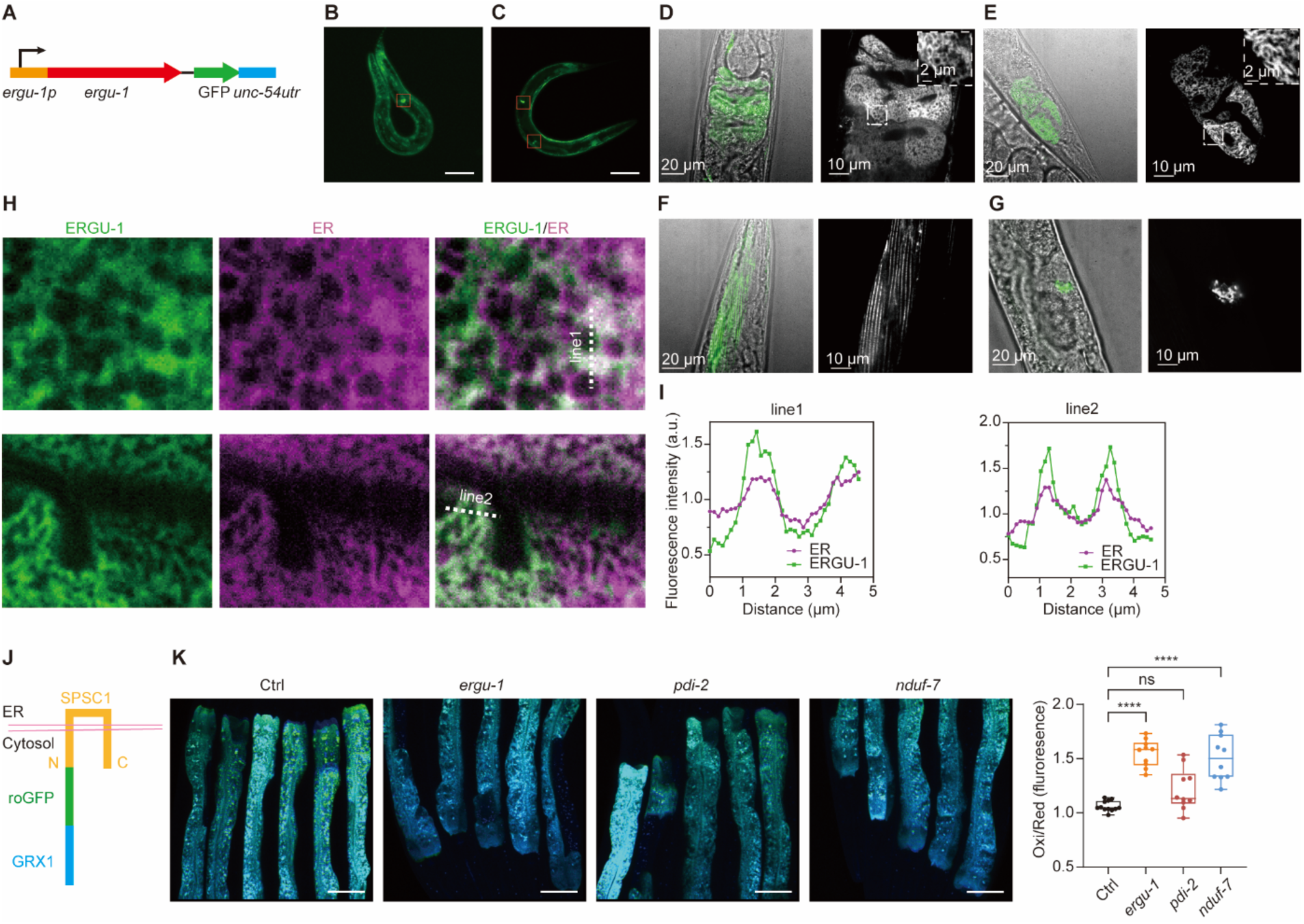
ERGU-1 constitutes a net-like reticulum structure at ER membranes. (**A**) Schematic of the *ergu-1*p::ERGU-1::GFP translation reporter. (**B** and **C**) Representative low-mag view of *ergu-1*p::GFP (B) and *ergu-1*p::*ergu-1*::GFP (C) respectively, showing expression in anterior and posterior intestine, body wall muscles and spermatheca. Scale bars, 100 μm. (**D** to **G**) Representative high-mag confocal view of *ergu-1*p::ERGU-1::GFP showing protein localization and expression in anterior (D) and posterior (E) intestinal cells, body wall muscles (F) and spermatheca (G). (**H**) Representative high-mag confocal view of ERGU-1::GFP proteins showing co-localization with the ER membrane marker SEL-1(1-79)::mCherry::HDEL in the intestine, body wall muscles and spermatheca. (**I**) Line scans showing co-localization. Scale bars, 100 μm. (**J**) Schematic design of the ER-proximal H_2_O_2_ sensor GRX1-roGFP-SPSC-1. (**K**) Representative ratio metric images and quantification of elevated H_2_O_2_ levels in the cytosol near the ER under *ergu-1* and *nduf-7* RNAi treatment, as indicated by the ER-proximal H_2_O_2_ sensor. Scale bars, 100 μm.

To ascertain ERGU-1’s subcellular localization, we crossed the translational reporter with a previously well-characterized ER membrane marker SEL-1(1-79)::mCherry::HDEL^28,29^. High-resolution dual-fluorescence confocal microscopy and line-scanning analysis revealed marked co-localization of *ergu-1*p::*ergu-1*::GFP and SEL-1(1-79)::mCherry::HDEL in a net structure-like reticulum pattern (Fig. 3D-3I, Fig. S5A). ERGU-1::GFP showed particularly prominent intensities in anterior and posterior intestinal cells, consistent with effects we observed with roGFP2-Orp1 (Fig. 2B). We confirmed the ER membrane-specific localization of ERGU-1 by imaging ERGU-1::GFP alongside RFP markers specific for other organelles, including the mitochondria, lysosomes, and plasma membrane (Fig. S5B). The ER membrane localization aligns markedly well with ERGU-1’s proposed functional role in mitigating the detrimental effects of H_2_O_2_ produced during protein folding within the ER lumen. To address potential ER membrane-localized changes in oxidative stresses, we generated ER-proximal redox sensor strain GRX1-roGFP-SPSC-1. We observed markedly higher oxidation to reduction ratio of roGFP upon *ergu-1* and *nduf-7* (positive control) RNAi treatment (Fig. 3J, 3K). Furthermore, biochemical purification of *ergu-1*p::*ergu-1*::GFP using GFP-trap affinity chromatography revealed a redox-sensitive oligomeric organization of ERGU-1 (Fig. 4F), with monomers being prevalent under reducing conditions. The ER membrane-specific localization, redox regulation and antioxidant function support ERGU-1 as a guardian against peroxide stress originating from ER.

**Fig. 4.**
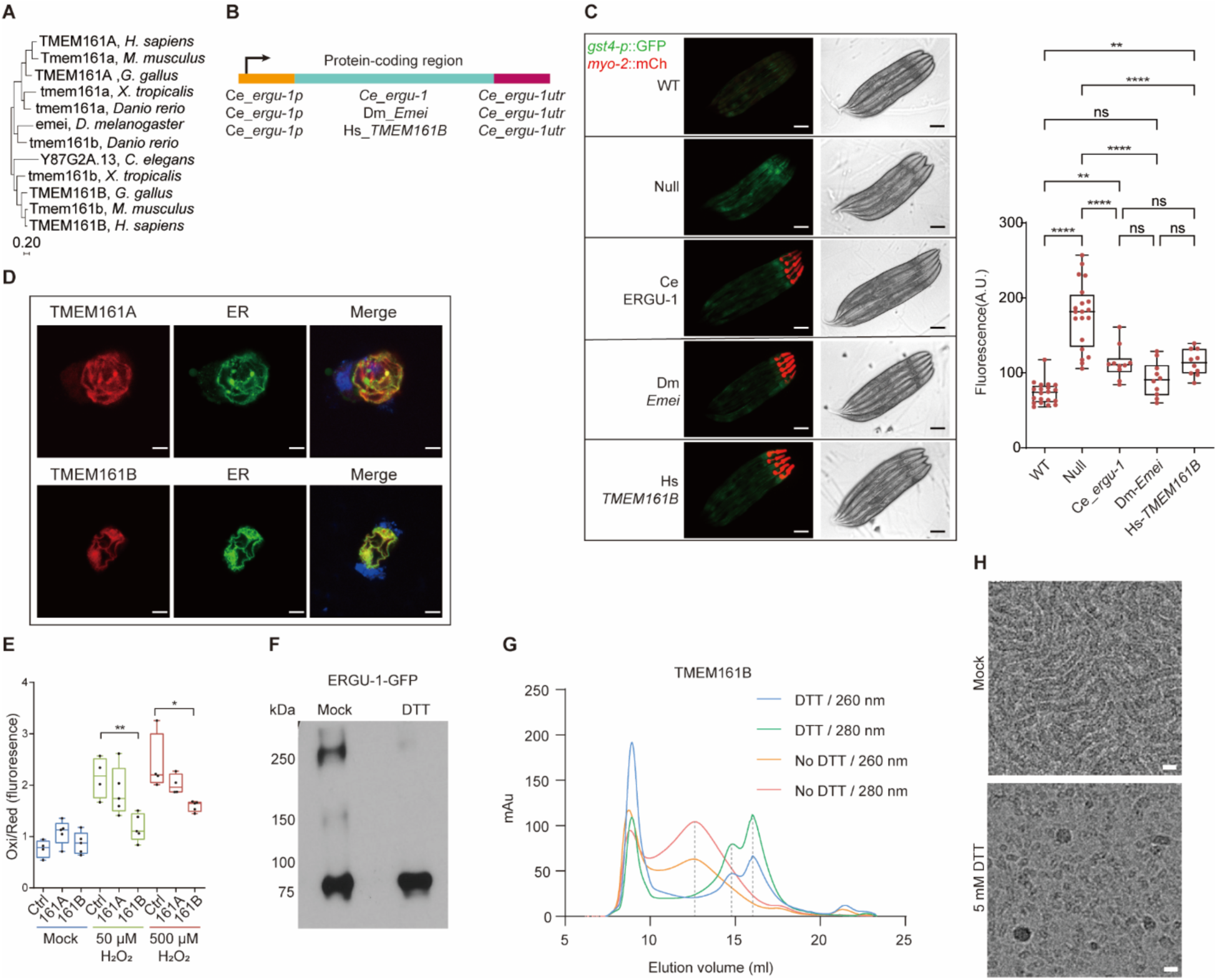
Conserved ER localization and antioxidant roles of ERGU-1 homologs. (**A**) Maximum likelihood-based phylogenetic tree of ERGU-1 family proteins. (**B**) Schematic representation of transgenic constructs and strategy to rescue *ergu-1* null phenotype. (**C**) Representative fluorescence images of *gst-4p::GFP* showing rescue of *C. elegans ergu-1* mutants by *Drosophila* and human homologs (n=10 for each rescue group). Quantification (right). Scale bars, 100 μm. (**D**) Immunofluorescence images showing that Human TMEM161A/B localizes to the ER membrane in 293T cells. Scale bars, 10 μm. (**E**) Reduction of cytosolic H_2_O_2_ detected by roGFP2-Orp1 in 293T cells overexpressing human TMEM161A or TMEM161B. (**F**) Representative SDS-PAGE western blots with antibodies against GFP showing formation of oligomers (likely dimers and tetramers based on the molecular weight) by ERGU-1::GFP and reduction to monomers by 10 mM dithiothreitol (DTT). (**G**) Chromatography plot showing reduction of purified human TMEM161B to smaller size peaks in the presence of 5 mM DTT as analyzed by SEC. (**H**) Reduction of TMEM161B strep-HA from filamentary forms to monomers or dimers in 5 mM DTT as shown by cryo-electron microscopy. Scale bars, 10 nm. [(C) and (E)] *P* value was determined by one-way ANOVA.

### Evolutionarily conserved features of ERGU-1 family proteins

We next investigated the evolutionary conservation of ERGU-1. First, we constructed a multiple sequence alignment and a maximum likelihood-based phylogenetic tree encompassing ERGU-1 and its homologs from diverse animal species (Fig. 4A). This analysis revealed a high degree of sequence conservation across the ERGU-1 protein family, suggesting a potentially ancient and functionally important role. For instance, the mouse ERGU-1 homolog Tmem161b and human TMEM161B exhibit 93.6% amino acid percent identities, while *C. elegans* ERGU-1 and human TMEM161B exhibit 27.9% amino acid percent identities. To experimentally test evolutionary conservation of the ERGU-1 protein family, we sought to functionally complement the *C. elegans ergu-1* mutant phenotype with homologs from other species. We expressed the *ergu-1* homologs *emei* and *TMEM161B* from *Drosophila* and humans, respectively, under the control of endogenous *C. elegans ergu-1* promoter, in the *ergu-1* mutant background carrying the *gst-4p::GFP* reporter (Fig. 4B). Remarkably, we found that transgenic expression of these homologs rescued the constitutively activated *gst-4p::GFP* phenotypes, indicating normalized oxidative stress in *C. elegans ergu-1* mutants (Fig. 4C). This successful complementation across species supports the functional conservation of ERGU-1 and its homologs within the ER-GUARD system.

To additionally explore the functional conservation of ERGU-1 homologs, we focused on TMEM161B, which appears more closely related to ERGU-1 than its paralog TMEM161A (Fig. 4A). To mirror our *C. elegans* studies, we first examined TMEM161B’s subcellular localization in mammalian cells. We expressed V5-tagged TMEM161B cDNA under the control of a CMV promoter in HEK293T and U2OS human osteosarcoma cells. Immunofluorescence staining using an anti-V5 antibody revealed a striking colocalization with established ER organelle markers (Fig. 4D, Fig. S6F). Furthermore, to assess the potential antioxidant function of TMEM161B as *C. elegans* ERGU-1, we employed V5-tagged TMEM161B cDNA expressed in HEK293 human embryonic kidney cells. With the H_2_O_2_ sensor roGFP2-Orp1, we found that cDNA expression of human TMEM161B led to strong suppression of exogenous H_2_O_2_-induced oxidation of roGFP2-Orp1 (Fig. 4E, Fig. S6G). We also used the latest AlphaFold3 webserver^30^ to model various ligand interaction with TMEM161B and ERGU-1, and identified heme-binding pockets coordinated by a highly conserved tyrosine residue (Y454) (Fig. S4B). We experimentally confirmed that biochemically purified TMEM161B, but not Y454 mutants, bound to heme directly, while functionally the Y454 mutant ERGU-1 or TMEM161B failed to rescue *ergu-1* null mutants (Fig. S4C). Beyond the redox-sensitive oligomeric organization of ERGU-1 (Fig. 4F), purified human TMEM161B exhibited native oligomerization under normal conditions and dissociation into monomers in a reducing environment, as demonstrated by size exclusion chromatography (Fig. 4G). Interestingly, cryo-electron microscopy analysis revealed a filamentary oligomeric form of TMEM161B under non-reducing conditions, while treatment with DTT induced monomerization (Fig. 4H). These findings indicate that human TMEM161B recapitulates several key features of *C. elegans* ERGU-1, including ER membrane localization, antioxidant functions and redox-modulated oligomeric states, providing further evidence for the evolutionary conservation of ERGU-1 in antioxidant defense across the animal kingdom.

## Discussion

In this study, we identify ERGU-1, a previously uncharacterized and evolutionarily conserved protein, in a critical ER-resident antioxidant defense system in animal cells. We used a combination of computational and functional screening to pinpoint ERGU-1 among the entire UniProt-defined *C. elegans* proteome. Determining the initial list of ERGU candidates for functional validation exemplifies the power of AlphaFold2, an AI for protein structure prediction. Notwithstanding known caveats of predicted structures by AlphaFold2 as compared to experimentally determined ones in accuracy, predicted structures can guide researchers towards promising avenues for functional studies, leading to advances in our understanding of fundamental biological processes and potentially paving the way for new therapeutic strategies^30–33^. We designed the filtering criteria for screening ERGU candidates based on empirical knowledge gained from DsbD/ScsB and membrane oxidoreductases. Similar strategy based on modified criteria could lead to discovering other protein functions beyond ERGU-1. More broadly, this approach also highlights the potential of AI in accelerating new biological discovery.

Despite the known generation of H_2_O_2_ during oxidative protein folding, the identity of the ER membrane-resident protein responsible for its detoxification has remained elusive. Our work bridges this gap by identifying ERGU-1 as such a key player, which harbors functional and structural features characteristic of transmembrane oxidoreductases. Importantly, genetic deletion of ERGU-1 led to constitutive activation of oxidative stress- and ER stress-response reporters, elevated cytosolic H_2_O_2_ levels, and organismal phenotypes in reproduction and behaviors. These findings strongly suggest ERGU-1’s crucial role in mitigating ER-derived oxidative stress, and mechanistically by transferring electrons via cysteine pairs across ER membranes, with ER luminal H_2_O_2_ as a likely electron acceptor. As loss of ERGU-1 also activated the *hsp-4p::GFP* reporter indicative of unfolded protein stress, it is plausible that ERGU-1 provides reducing equivalents from cytosolic electron donors for additional ER substrates other than H_2_O_2_, e.g. misfolded proteins and certain peroxide products that ScsB or DscB can act upon to reduce^12,34,35^. However, it is worth noting that neither ScsB nor DsbD is known to bind to heme, unlike the *C. elegans* and human homologs of ERGU-1. We thus postulate that eukaryotic ER-GUARD may have evolved additional heme-binding capacity that likely enables efficient electron transfer together with cysteine pairs.

The subcellular localization of ERGU-1 at the ER membrane is well suited for its antioxidant function. By residing at the site close to H_2_O_2_ generation, ERGU-1 may efficiently neutralize it, safeguarding the ER membrane and nearby cytosol from oxidative damage. Furthermore, the redox-sensitive nature of the tetrameric ERGU-1 organization suggests a potential regulatory mechanism for ER-GUARD function. Under more oxidizing conditions, the tetrameric form appears to be more prevalent, potentially enhancing ERGU-1’s antioxidant activity to safeguard against excess peroxide stress. Interestingly, our findings on ER-GUARD and its proximity to H_2_O_2_ generation in the ER align with recent studies suggesting a limited role for mitochondrial ROS in causing global oxidative stress and nuclear DNA damage^36^. This highlights the ER as a potential major source of cellular oxidative stress impacting nuclear membranes and the genome. By effectively neutralizing ER-derived H_2_O_2_, ERGU-1 may play a critical role in shielding the genome from oxidative damage. Furthermore, ribosomal RNA, densely packed on ER membranes and crucial for protein synthesis, is another likely target of such oxidative damage. It remains determined whether ERGU-1 protects against oxidative damage in general or more specifically biases towards certain biomolecules. In specific tissues (e.g. intestine), excess peroxide stress caused by loss of ERGU-1 may also activate multiple compensatory antioxidant pathways, including SKN-1/NRF2, and recruit additional mechanisms to safeguard cellular and organismal functions.

The high degree of sequence conservation observed within the ERGU-1 protein family across diverse animal species underscores its potential evolutionary ancestry and functional importance. The successful complementation of the *C. elegans ergu-1* mutant phenotype with homologs from *Drosophila* and humans further strengthens this notion. Although we did not find any apparent orthologues of ERGU-1 in eukaryotic fungi, our findings indicate evolutionary conservation of the ER-GUARD across the animal kingdom, emphasizing its critical and previously undescribed role in maintaining cellular redox balance. Mutations in the *Drosophila* orthologue *emei* impair ER calcium dynamics^37^, whereas mutations in its vertebrate orthologue *Tmem161B* cause severe pathological cardiac arrhythmias in mice and brain polymicrogyria in humans^38–41^. Our studies suggest that these previously unexplained phenotypic defects might be mechanistically caused by defects in ER-GUARD functions and redox homeostasis. Thus, future studies are warranted to investigate the mechanisms of action, redox regulation of ER-GUARD, and its functional consequences at the cellular and organismal levels that may be broadly important for human physiology and diseases.

## Supporting information

Python codes for AlphaFold screens

---

**Fig. S1.**
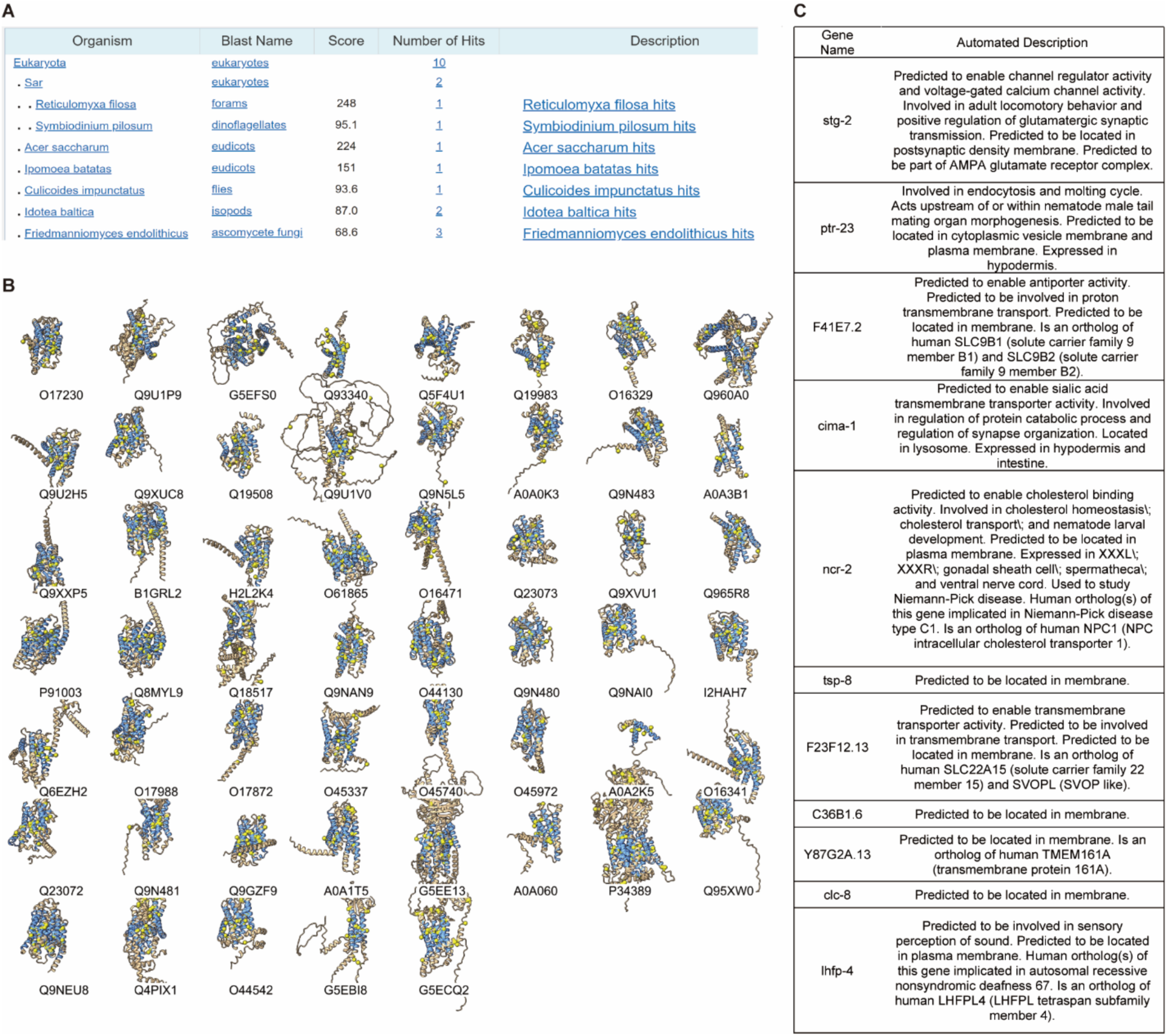
AlphaFold2-based computational screen for ERGU candidates. (**A**) BLASTP search yielding no apparent broadly conserved eukaryotic homologs of DscB/ScsB based on protein sequence similarities, prompting a search for DscB/ScsB homologs based on functional and structural features using AlphaFold2-predicted structures. (**B**) Predicted structures of the 53 ERGU candidates identified. (**C**) Table listing the annotated functions of the 11 non-GPCR ERGU candidates.

**Fig. S2.**
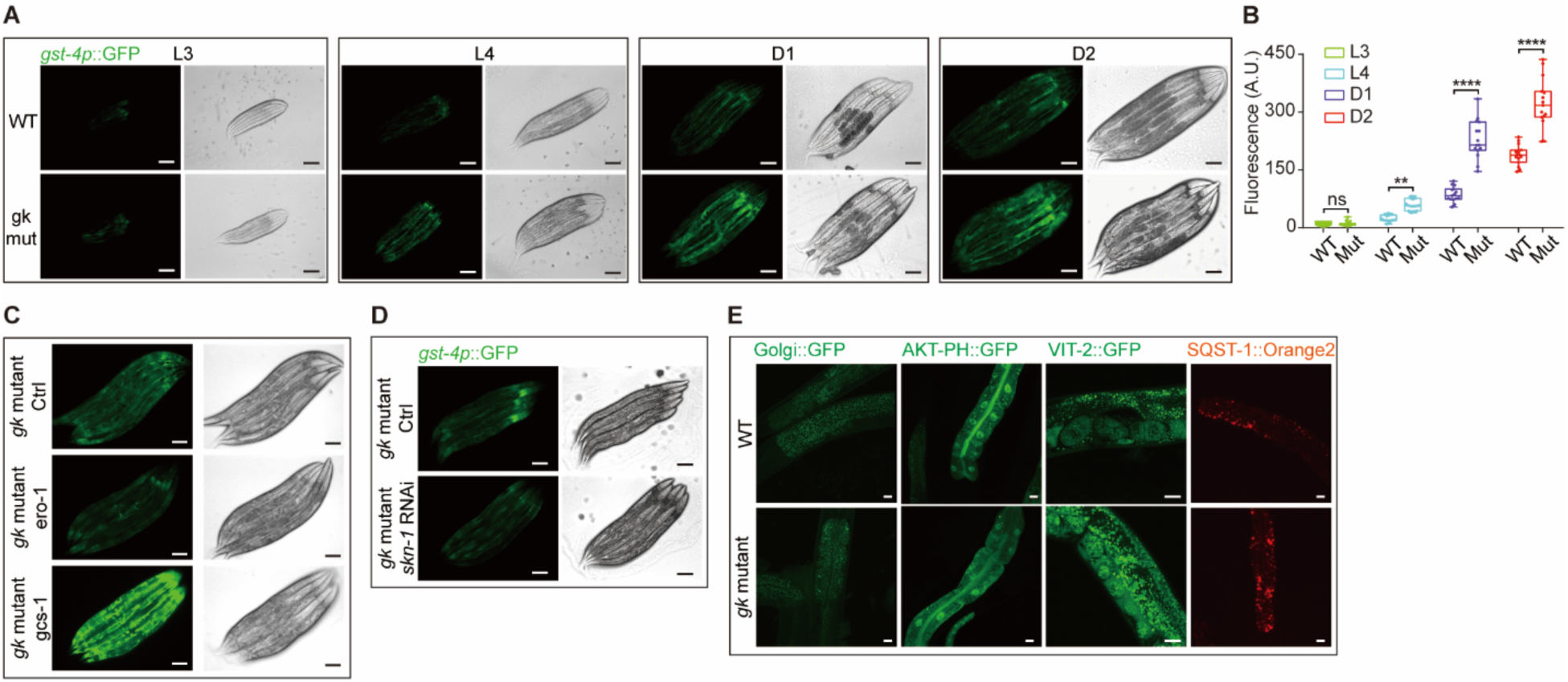
Phenotypes of ERGU-1 protein-truncating *ergu-1(gk840471)* mutants. (**A**) Representative epifluorescence images showing *gst-4p::GFP* activation in *ergu-1*(*gk840471*) mutants (n=15 for each group). (**B**) Quantification of *gst-4p*::GFP fluorescence intensities in WT and *ergu-1*(*gk840471*) mutants indicated (n=15 for each group). (**C**) Representative epifluorescence images showing *gst-4p::*GFP decrease by *ero-1* RNAi and increase by *gcs-1* RNAi (**D**) Representative epifluorescence images showing *gst-4p::GFP* activation in *ergu-1*(*gk840471*) mutants requires *skn-1.* (**E**) Representative confocal fluorescence images showing no apparent reporter (Golgi apparatus-marking Golgi::GFP, apical and endosomal membrane-marking AKT-PH::GFP, yolk organelle-marking VIT-2::GFP and autophagosome-marking SQST-1::mOrange2) alteration in *ergu-1*(*gk840471*) mutants. *P* value was determined by an unpaired *t*-test, two-tailed (comparison between two groups). [(A), (B) and (C)] Scale bars, 100 μm. (D) Scale bars, 20 μm.

**Fig. S3.**
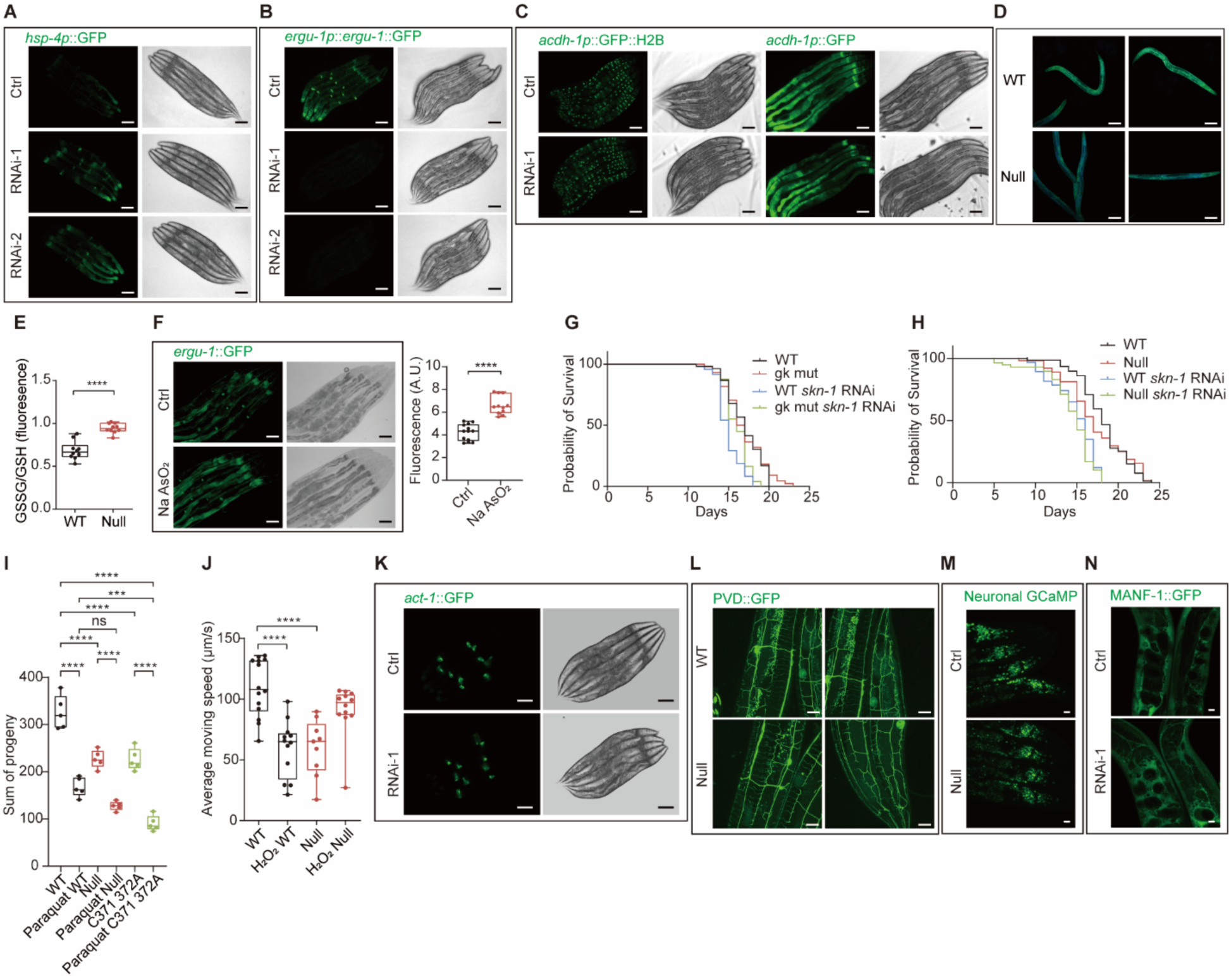
Phenotypic consequences of RNAi or mutant *ergu-1*. (**A**) Representative epifluorescence images showing increased *hsp-4p::GFP* upon *ergu-1* RNAi treatment. (**B**) Representative fluorescence images of *ergu-1p*::*ergu-1::GFP* showing abolished GFP signals by *ergu-1* RNAi. (**C**) Representative fluorescence images of Vitamin B12 stress reporter *acdh-1p*:: *GFP* showing unaltered GFP signals by *ergu-1* RNAi. (**D** and **E**) Representative fluorescence images of glutathione stress GSSG/GSH reporters showing elevated oxidative stress in *ergu-1* null mutants (D) and quantification (n=10) (E). (**F**) Representative fluorescence images of *ergu-1p*::*ergu-1::GFP* showing reporter up-regulation by oxidative stressors (0.5% NaAsO_2_). (**G**) Lifespan curves of wild-type and *ergu-1* null mutants with control or *skn-1* RNAi showing comparable lifespans in wild-type and *ergu-1* null mutants and reduction by *skn-1* RNAi. (**H**) Lifespan curves of wild-type and *ergu-1* protein-truncating mutants (*gk840471*) with control or *skn-1* RNAi comparable lifespans in wild-type and *ergu-1(gk840471)* mutants and reduction by *skn-1* RNAi (n ≥ 50). (I) Quantification of brood sizes in wild type, *ergu-1* null and cysteine mutants upon 10 mM paraquat treatment. *P* value was determined by one-way ANOVA (comparison between multiple groups). (**J**) Quantification of average moving speed in wild-type, ergu-1 null and above strain treated with 10 mM H2O2. (**K**) Representative fluorescence images of cytoskeletal actin ACT-1::GFP reporters showing no apparent alteration by *ergu-1* RNAi. (**L**) Representative confocal fluorescence images showing no apparent reporter (PVD neuron-marking GFP) alteration in *ergu-1* null mutants. (**M**) Representative confocal fluorescence images showing no apparent reporter (neuronal nuclear GCaMP6) alteration in *ergu-1* null mutants. (**N**) Representative confocal fluorescence images showing no apparent reporter (secreted MANF-1::GFP) alteration by *ergu-1* RNAi. *P* value was determined by an unpaired *t*-test, two-tailed (comparison between two groups). [(A) to (J)] Scale bars, 100 μm. [(K) to (M)] Scale bars, 20 μm.

**Fig. S4.**
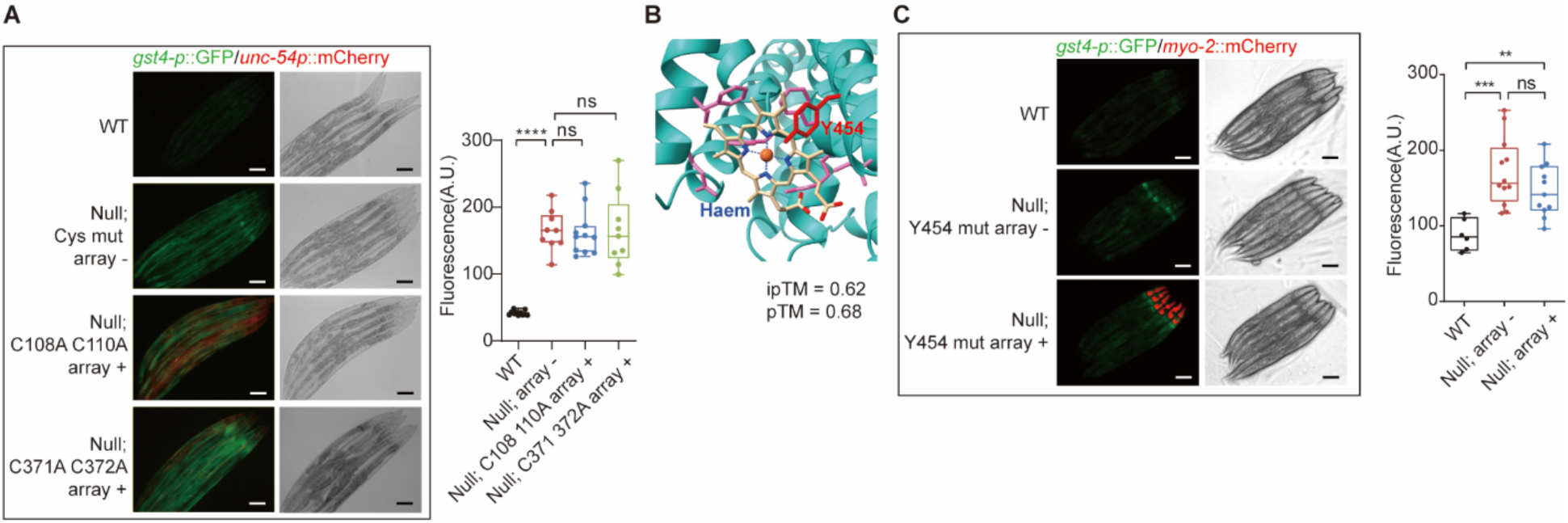
Importance of key cysteine residues for ERGU-1 functions. (**A**) Representative fluorescence images of *gst-4p::GFP* showing no rescue of *C. elegans ergu-1* mutants by two different cysteine pair mutants of ERGU-1. (**B**) Schematic of key residues for heme binding and interactions. Heme binding pocket is in high confidence region (90 > plDDT > 70). (**C**) Representative fluorescence images and quantification of *gst-4p::GFP* showing no rescue of *C. elegans ergu-1* mutants by *C. elegans* or human Y454 mutants of ERGU-1 homologs. [(A) and (C)] Scale bars, 100 μm.

**Fig. S5.**
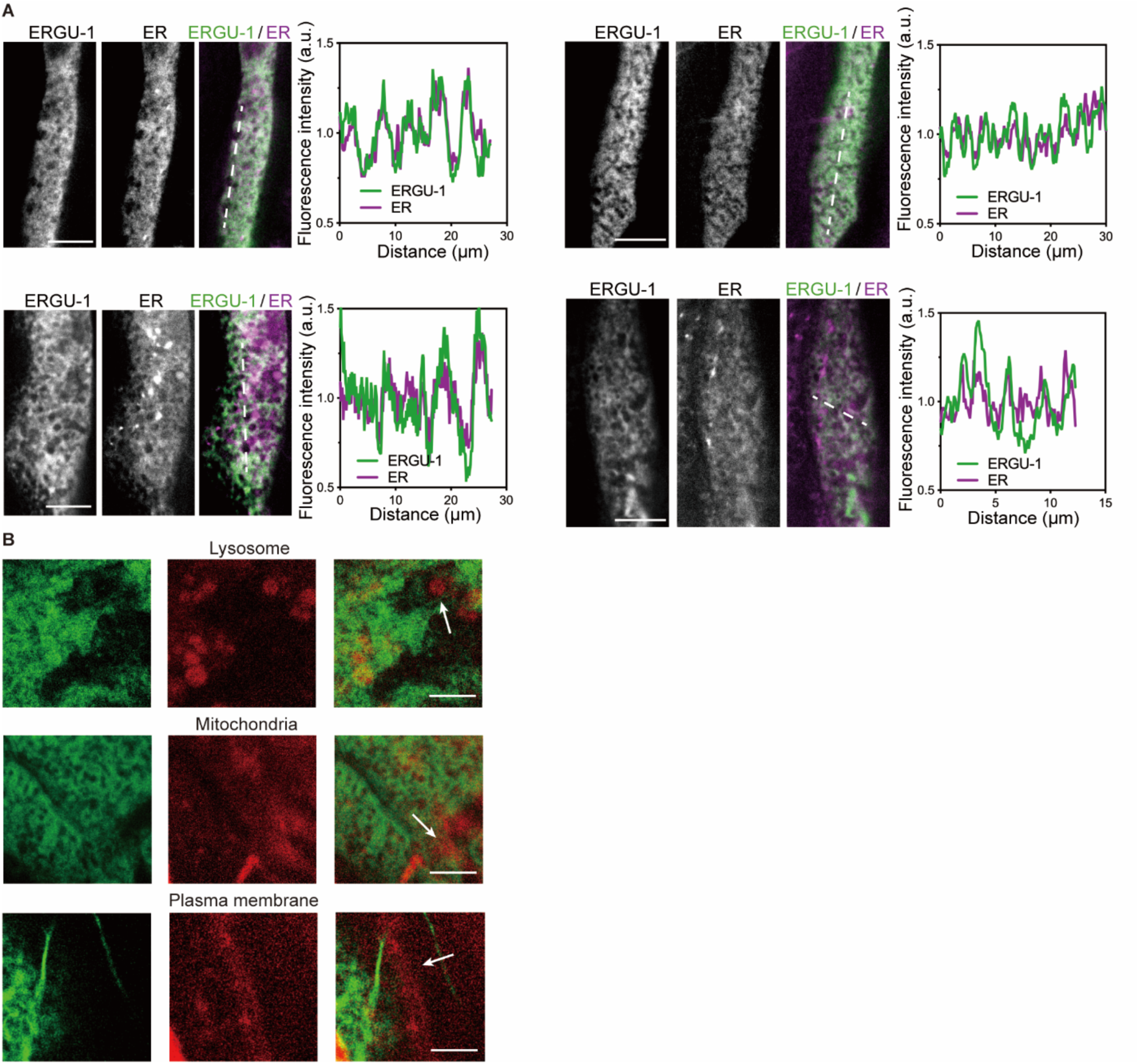
Colocalization of ERGU-1::GFP with RFP-marked ER organelle reporters but not non-ER organelle reporters. **(A)** Representative confocal fluorescence images showing colocalization between ERGU-1::GFP (green) and endoplasmic reticulum-specific reporters (red). Scale bars, 10 μm. (**B**) Representative confocal fluorescence images showing no colocalization between ERGU-1::GFP (green) and lysosome, mitochondria or plasma membrane-specific RFP reporters (red). Arrow indicates the organelles. Scale bars, 5 μm.

**Fig. S6.**
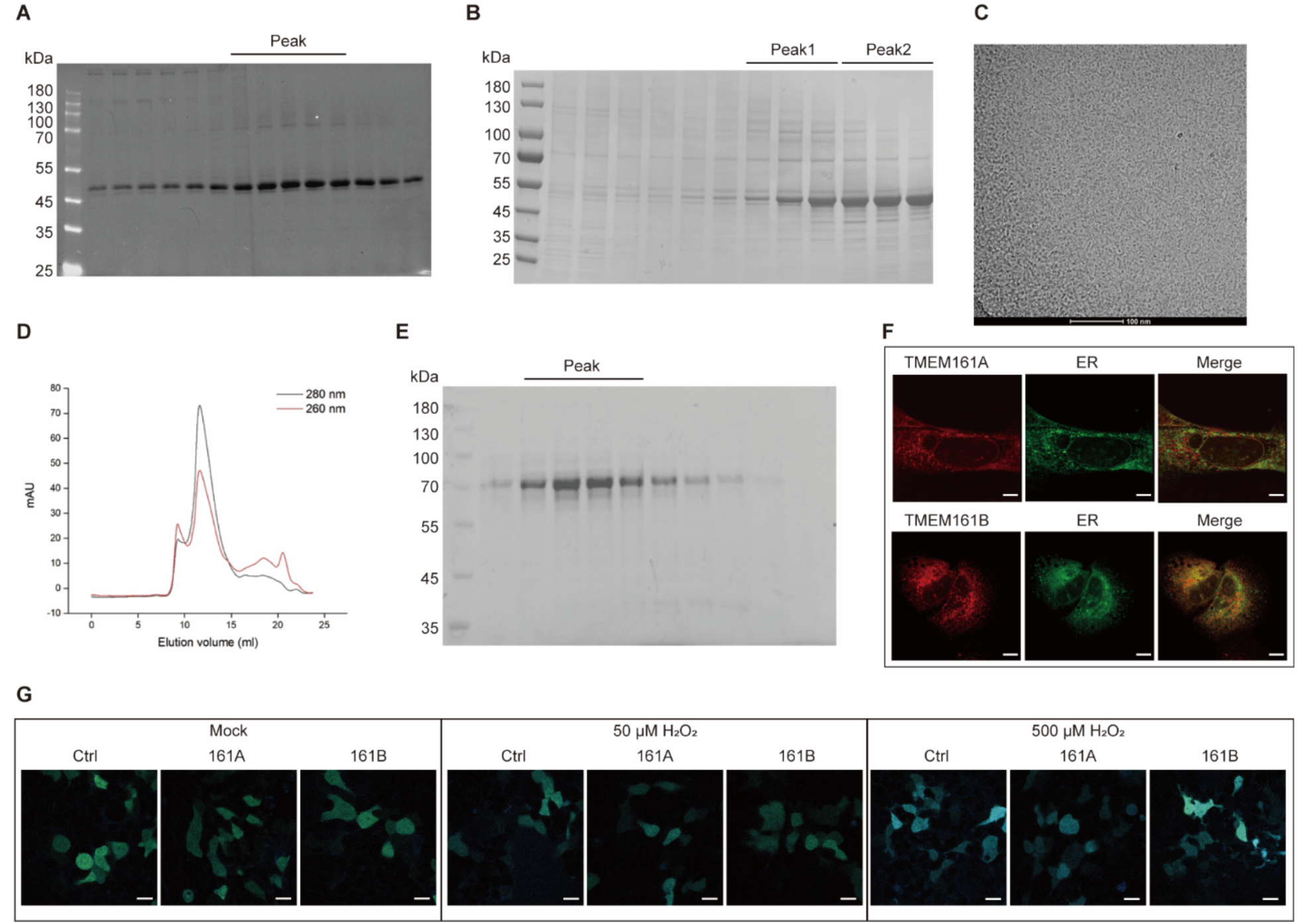
Protein characteristics of human TMEM161B, an ERGU-1 ortholog and evolutionarily conserved member of the ERGU protein family. (**A**) Peak fractions of human TMEM161B-2X Strep-HA protein purified by SEC in a non-reductive environment. (**B**) Peak fractions of TMEM161B-2X Strep-HA protein purified by SEC in a DTT-containing reductive environment. (**C**) TMEM161B-GFP in a non-reductive environment shown by Cryo-EM. (**D**) Peak of TMEM161B-GFP in a non-reductive environment shown by SEC. (**E**) Peak fractions of TMEM161B-GFP protein purified by SEC in a non-reductive environment. (**F**) Immunofluorescence images showing that Human TMEM161A/B localizes to the ER membrane in U2OS cells. Scale bars, 10 μm. (**G**) Representative confocal fluorescence images of roGFP2-Orp1 detecting cytosolic H_2_O_2_ in 293T cells overexpressing human TMEM161A or B. Scale bars, 20 μm.

## Materials and Methods

### C. elegans

*C. elegans* strains were maintained under standard laboratory conditions unless otherwise specified. The N2 Bristol strain was used as the reference wild-type. Feeding RNAi was performed as previously described^42^. Transgenic strains were generated by germline transformation as described^43^. Transgenic constructs were co-injected (at 10 - 50 ng/μl) with dominant *unc-54p*::mCherry or *myo-2p*::mCherry, and stable extrachromosomal lines of fluorescent animals were established for UV-mediated genome integration. Genotypes of strains used are as follows: *ergu-1(ust572, syb9153, syb9160), dvIs19 [gst-4p::GFP], zcIs4 [hsp-4::GFP], jrIs10 [unc119(+) rps-0p::roGFP2-Orp1], dmaIs160 [ergu-1p:: ergu-1::gfp; unc-54p::mCherry], dmaEx701[ergu-1p::hsTMEM161B::ergu-1utr; myo-2p::mCherry], dmaEx702 [ergu-1p::ergu-1::ergu-1utr; myo-2p::mCherry], dmaEx704 [ergu-1p::Dm-161:: ergu-1utr; myo-2p::mCherry]*.

### AlphaFold-assisted computational screen

A comprehensive computational screen of the *Caenorhabditis elegans* proteome was performed to identify potential DsbD/ScsB-like proteins. The reference proteome utilized for this analysis was the *C. elegans* UniProt reference proteome (version 26), encompassing 19,827 genes. Initial filtering was conducted to select genes encoding proteins with a minimum of four annotated transmembrane helices, resulting in a subset of 3,177 genes. Subsequent filtering criteria focused on the presence of transmembrane helices containing cysteine residues, specifically targeting helices with at least two cysteines separated by no more than two intervening residues, corresponding to the sequence motifs CC, CXC, or CXXC. This filtering narrowed the candidate list to 190 genes. The structural analysis was conducted using predicted protein structures from the AlphaFold2 Database. For each of the 190 genes, the predicted structures were examined to determine the spatial proximity of cysteine pairs located within the transmembrane helices. Specifically, structures in which the sulfur atoms (SG) of cysteine pairs were within 10 angstroms of each other were identified. This criterion was met by 53 candidate genes. All computational and structural analyses were executed using custom Python scripts, which are provided in the Supplementary file. Visualization and further structural analyses were performed using UCSF ChimeraX software.

### RNA interference (RNAi) cloning and screen

RNAi and screen for hits upregulating *gst-4p*::GFP were performed by feeding worms with E. coli strain HT115 (DE3) expressing double-strand RNA (dsRNA) targeting endogenous genes. Briefly, dsRNA–expressing bacteria were replicated from the Ahringer library to LB plates containing 100 μg/ml ampicillin (BP1760-25, Fisher Scientific) at 37 °C for 16 hrs. Single clone was picked to LB medium containing 100 μg/ml ampicillin at 37 °C for 16 hrs and positive clones (verified by bacteria PCR with pL4440 forward and pL4440 reverse primers) were spread onto NGM plates containing 100 μg/ml ampicillin and 1 mM isopropyl β-D-1-thiogalactopyranoside (IPTG, 420322, Millopore) for 24 hrs. Developmentally synchronized embryos from bleaching of gravid adult hermaphrodites were plated on RNAi plates and grown at 20 °C. The worms are collected for imaging at indicated stages.

To make the *ergu-1* RNAi-1 and RNAi-2 plasmids, RNAi-1 targeting sequences are PCR-amplified with primers 5’- GGCCCCCCCTCGAGGTCGACAGGAGACGATGCTATAAACTATGCT -3’ and 5’- TCCACCGCGGTGGCGGCCGCCGTGACGTGTTTTCAGCTCG -3’ from N2 gDNA and subcloned into the SalI and NotI sites of a digested T777T vector with Promega T4 DNA ligase (Promega, M1801). RNAi-2 targeting sequences are PCR-amplified with primers 5’-GGCCCCCCCTCGAGGTCGACATGCCAAAATGAATGCGGGC -3’ and 5’-TCCACCGCGGTGGCGGCCGCACGTGGTGATATGCCTGGTG -3’.

### Fluorescence microscopy and H_2_O_2_ sensor imaging

SPE confocal (Leica) and epifluorescence microscopes were used to capture fluorescence images. Animals were randomly picked at the same stage and treated with 1 mM levamisole in M9 solution (31742-250MG, Sigma-Aldrich), aligned on a 2% agar pad on a slide for imaging. Identical setting and conditions were used to compare experimental groups with control. For quantification of GFP fluorescence, animals were outlined and quantified by measuring gray values using the ImageJ software. The data were plotted and analyzed by using GraphPad Prism10. For *jrIs10 [unc119(+) rps-0p::roGFP2-Orp1]* strain, orp1-roGFP2 was excited sequentially at 405 and 488 nm and emission was recorded at 500-540 nm. Fifteen images were sequentially captured at 1-micrometer z-intervals and subsequently stacked to form a composite image.

### Western blotting

For SDS-PAGE samples, stage-synchronized animals for control and experiment groups were picked (n = 50) in 60 μl M9 buffer and lysed directly by adding 20 μl of 4x Laemmli sample buffer (1610747, Bio-Rad) contain 10% of 2-Mercaptoethanol (M6250-100ML, Sigma (v/v)). Protein extracts were denatured at 95 °C for 10 min and separated on 10% SDS-PAGE gels (1610156, Bio-Rad) at 80 V for ∼40 min followed by 110 V for ∼70 min. The proteins were transferred to a nitrocellulose membrane (1620094, Bio-Rad,) at 25 V for 45 mins by Trans-Blot® Turbo™ Transfer System (Bio-Rad). The NC membrane was initially blocked with 5% non-fat milk and 2% BSA (A4503, Sigma (v/v)) in tris buffered saline with 0.1% Tween 20 (93773, Sigma) (TBST) at room temperature for 1 h. Proteins of interest were detected using antibodies against GFP (A6455, Invitrogen) or V5 (13202T, Cell Signaling Technology) in cold room for overnight. After three washes of 5 min each with tris-buffered saline with 0.1% Tween 20, anti-mouse IgG, HRP-linked Antibody (7076, Cell Signaling Technology) was added at a dilution of 1:5000. For DTT treatment, worm lysates were treated with 10 mM DTT and boiled at 95 °C for 10 min.

### Mammalian cell culture experiments

Human embryonic kidney (HEK) 293T cells and osteosarcoma U2OS cells were maintained in Dulbecco’s modified Eagle’s medium with 10% inactive fetal bovine serum and penicillin-streptomycin (Gibco, Grand Island, 15140122) at 37 °C supplied with 5% CO2 in an incubator (Thermo Fisher Scientific) with a humidified atmosphere. Cells were washed once using PBS and digested with 0.05% trypsin EDTA (Gibco) at 37 °C for routine passage of the cells. All cells were transfected with 3 μl LipoD293^TM^ (SignaGen, SL100668) per 1 μg DNA mixture. HEK293T cells were transfected by the pLX304-TMEM161A-V5 (DNASU, HsCD00441633) or pLX304-TMEM161B-V5 (DNASU, HsCD00444935) and collected 2 days after transfection. HEK293T and U2OS cells were transfected by the pHAGE2-TMEM161A-gfp/mCherry or pHAGE2-TMEM161B-gfp/mCherry and collected for imaging 2 days after transfection. pHAGE2-TMEM161A/B-GFP/mCherry was constructed by inserting TMEM161A or TMEM161B into pHAGE2-gfp/mCherry plasmids. TMEM161A was PCR-amplified with the primers 5’-TAATTAAACCTCTAGAGCCACCATGGCGGTCCTCGG -3’ and 5’-CTCACCATAGCTCGAGacccagacccGGAGCCTGCCAAGTGC -3’ from pLX304-TMEM161A-V5 mentioned before. TMEM161B was PCR-amplified with the primers 5’-TAATTAAACCTCTAGAGCCACCATGGGTGTGATAGGTATACAGC -3’ and 5’-CTCACCATAGCTCGAGacccagacccTGCCACAGTCAGATACTGGT -3’ from pLX304-TMEM161B-V5 mentioned before.

### Quantification of brood size

The brood size assay was carried out according to the standard protocol. (1) Briefly, single L4 worms of different stains (N2, ergu-1 null, PHX9155, PHX9160) were individually placed in 60mm Petri plates and kept in the incubator at 20 C. Then, the worms were transferred to a new OP50 containing plate each day at the same time till day 6 of the worms. Progenies were scored after 3 days and plotted for each day and the total sum of progenies. For NAC treatment the plates were supplemented with 10 mM of NAC during the entire duration of the assay. For H_2_O_2_ and paraquat treatment the plates were supplemented with 10 mM of H_2_O_2_ and paraquat respectively during the assay. Each experiment was repeated 3 times as independent biological replicates with more than 5 animals per group.

### Behavioral assay

D1 worms (24 hrs post L4) were transferred to a fresh NGM plate seeded with a small and thin OP50 bacterial lawn and allowed to settle for at least ten minutes to recover at room temperature. Moving average speeds, and track lengths of *C. elegans* were monitored for 10 minutes and were analyzed using WormLab. For H_2_O_2_ treatment, D1 worms are treated with 10 mM H_2_O_2_ for 30 min before assay.

### Lifespan analysis

For lifespan assays, Animals were cultured under non-starved conditions for at least 2 generations before life span assays. For normal NGM life span assay, stage-synchronized L4 stage animals (n ⩾ 50) were picked to new NGM plates seeded with OP50 containing 50 μM 5-fluoro-2′-deoxyuridine (FUDR) to prevent embryo growth at 20 °C incubator. Animals were scored for survival per 24 hrs. Animals failing to respond to repeated touch of a platinum wire were scored as dead. Three biological replicates were performed, with population sizes larger than 50 in each trial.

### DCFDA ROS detection

ROS staining in live worms was carried out as in standard protocol with minor modifications. Briefly, day 1 worms were washed three times with M9 and then transferred into 200 μl of M9 buffer containing 10 mM H2DCFDA (carboxy-H2DCFDA [5-(and-6)-carboxy-2′,7’-dichlorodihydrofluorescein diacetate) in an 1.5 ml Eppendorf tube and incubated in the dark for 3 hrs. The worms were randomly selected and treated with 10 mM sodium azide (Sigma-Aldrich) in M9, symmetrically aligned on 2% agar pads on slides for imaging the oxidized dichlorofluorescin (DCF).

### Expression and purification of TMEM161B for cryo-EM

The complementary DNA encoding human TMEM161B was cloned into a modified pDNA3.1 vector with a C-terminal twin-strep tag and a HA tag. For expression, Expi293F cells (Thermo Fisher Scientific) cultured in Freestyle293 Expression Medium were transfected with the vector DNA/polyethylenimine (1 μg DNA per ml culture, w/w = 1:3) complex at a cell density of ∼1.0 × 106 cells per ml and incubated at 37 °C under 8% CO2 with agitation at 100 r.p.m for 60 h. Cell pellets were resuspended in buffer A (25 mM Hepes pH 7.5, 150 mM NaCl, protease inhibitor cocktail) and were disrupted by sonication. For TMEM161B monomer purification, an extra incubation with 5 mM DTT at room temperature after sonication was required. The cell lysate was then spun at 150,000× g for 1h to sediment crude membranes. The membrane pellet was mechanically homogenized in buffer A. The suspension was solubilized in 1% (w/v) 1.0% DDM (Anatrance) and 0.1% CHS (Anatrance) for 60 min at 4 °C. The solubilized material was centrifuged at 100,000× g for 30 min, and the supernatant was incubated with Strep-Tactin®XT resins (iba) for 2 h at 4 °C. Resins were then washed with 20 column volumes of buffer B (25 mM Hepes pH 7.5, 0.025% DDM, 0.0025% CHS, 150 mM NaCl). The protein was eluted with buffer B supplied with 50 mM D-biotin, concentrated, and further purified by gel-filtration chromatography on a Superdex 200 increase column equilibrated with wash buffer B. The peak fractions of protein were pooled and concentrated to ∼5.0 mg/mL. For Cryo-EM observation, 3-µl aliquots of each sample were applied onto a glow-discharged Quantifoil grid (R1.2/R1.3 300 mesh, Au), blotted for 4.5–5.5 s in 100% humidity at 4 °C, and plunged into liquid ethane using a Vitrobot MkIV (Thermo Fisher Scientific). Cryo-EM micrographs were obtained by using a Talos Arctica G2 microscope (Thermo Fisher Scientific) running at 200 kV and a 300 kV Titan Krios microscope (Thermo Fisher Scientific).

### Statistical analysis

Data were analyzed using GraphPad Prism 9.2.0 Software (Graphpad, San Diego, CA) and presented as means ± S.D. unless otherwise specified, with *P* values calculated by unpaired two-tailed t-tests (comparisons between two groups), one-way ANOVA (comparisons across more than two groups) and adjusted with Bonferroni’s corrections. Box plots are presented as min to max, showing all data points.

## Acknowledgment

We are grateful for the feedback from and discussion with other members of the Ma lab at UCSF and Drs. P. Sigala, P. R. Ortiz de Montellano, A. Correia, Z. Chen, B. DeGrado, I. Jain, B. Black and G. Huang. We thank the *Caenorhabditis* Genetics Center (NIH grant #P40 OD010440), Wormbase.org (NIH grant #U24 HG002223 to P. Sternberg), Wormatlas.org (NIH grant #OD010943 to D.H. Hall.) and CenGen (cengen.org) for their immensely helpful resources. The work was supported by NIH grants (R35GM139618 to D.K.M., R35GM118167 to O.D.W), BARI Investigator Award (D.K.M.), and UCSF PBBR New Frontier Research Award (D.K.M., O.D.W).

## Author contributions

Z.J. and D.K.M. designed, performed and analyzed the *C. elegans* and cell culture experiments, contributed to project conceptualization and wrote the manuscript. T.P., B.W., Y.T. and S.H.G. designed, performed and analyzed the *C. elegans* transgenic and CRISPR experiments and editing manuscript. H.B. and Z.J. performed and analyzed the ER imaging experiments. T.D.G conducted the computational screen and contributed to project conceptualization and wrote the manuscript. Z.Y.L., J.X.Y., S.Y.X. contributed to human TMEM161B biochemical analysis. D.K.M, O.D.W. and T.D.G contributed to research materials, funding acquisition and editing manuscript.

