## Supplementary material for "AlphaFold2-Guided Functional Screens Reveal a Conserved Antioxidant Protein at ER Membranes": Python codes for AlphaFold screens

```
# Codes aim to find all C elegans proteins with pairs of close cysteines in transmembrane regions.

# Can use UniProt to identify transmembrane residues, then use AlphaFold database predicted structures

# to see if there are close cysteines.


def find_uniprot_transmembrane_cysteines(uniprot_xml_path, namespace = '{http://uniprot.org/uniprot}'):

    import xml.etree.ElementTree as ET

    tree = ET.parse(uniprot_xml_path)

    tm = []

    for child in tree.getroot():

        if child.tag == namespace + 'entry':

            rr = transmembrane_residue_ranges(child, namespace)

            uniprot_id = child.find(namespace + 'accession').text

            seq = child.find(namespace + 'sequence').text

            full_name = child.find(f'./{namespace}protein/{namespace}submittedName/{namespace}fullName')

            name = '' if full_name is None else full_name.text

            phc = paired_helix_cys(seq, rr)

            tm.append((uniprot_id, name, phc, rr))

    return tm


def paired_helix_cys(seq, rr):

    # Dengke suggests considering only paired cysteines CC, CxC or CxxC in a helix.

    phc = []

    for b,e in rr:

        ci = set([i+1 for i in range(b-1,e) if seq[i] == 'C'])

        pci = [i for i in ci

              if ((i+1) in ci or (i+2) in ci or (i+3) in ci or

                  (i-1) in ci or (i-2) in ci or (i-3) in ci)]
```

```
        if len(pci) >= 2:
            phc.append(pci)
    return phc

def transmembrane_residue_ranges(protein_xml_entry, namespace):
    ranges = []
    for feature in protein_xml_entry.iter(namespace + 'feature'):
        fattrib = feature.attrib
        if 'type' in fattrib and fattrib['type'] == 'transmembrane region':
            for loc in feature.iter(namespace + 'location'):
                b,e = loc.find(namespace + 'begin'), loc.find(namespace + 'end')
                if b is not None and e is not None:
                    if 'position' in b.attrib and 'position' in e.attrib:
                        r = (int(b.attrib['position']), int(e.attrib['position']))
                        ranges.append(r)
    return ranges

def close_cysteines(structure, membrane_residue_ranges, max_distance = 5):
    cys_res = [r for r in structure.residues if r.name == 'CYS']
    cys_xyz = [(r.number, r.find_atom('SG').coord) for r in cys_res]
    mb_res_nums = residue_numbers_from_ranges(membrane_residue_ranges)
    mb_cys = [r for r in cys_res if r.number in mb_res_nums]
    mb_xyz = [(r.number, r.find_atom('SG').coord) for r in mb_cys]

    close_pairs = set()
    from chimerax.geometry import distance
    for rnum, xyz in mb_xyz:
        for rnum2, xyz2 in cys_xyz:
            if rnum2 != rnum and distance(xyz, xyz2) <= max_distance:
```

```
            pair = (rnum, rnum2) if rnum < rnum2 else (rnum2, rnum)
            close_pairs.add(pair)

    return list(close_pairs)

def atoms_by_residue_number(atoms, atom_name):
    amap = {}
    for a in atoms:
        if a.name == atom_name:
            amap[a.residue.number] = a
    return amap

def residue_numbers_from_ranges(residue_ranges):
    res_nums = set()
    for b,e in residue_ranges:
        for rnum in range(b,e+1):
            res_nums.add(rnum)
    return res_nums

def check_for_close_cysteines(session, ulist, alphafold_dir, max_distance):
    found = []
    missing = []
    for uniprot_id, name, paired_hel_cys, tm_res_ranges in ulist:
        if len(paired_hel_cys) < 2:
            continue
        m = alphafold_database_model(session, uniprot_id, alphafold_dir)
        if m is None:
            missing.append((uniprot_id, name))
            continue
```

```
        close_pairs = []
        atoms = atoms_by_residue_number(m.atoms, 'SG')
        from chimerax.geometry import distance
        for i,ph1 in enumerate(paired_hel_cys):
            for ph2 in paired_hel_cys[i+1:]:
                for rnum1 in ph1:
                    for rnum2 in ph2:
                        if distance(atoms[rnum1].coord, atoms[rnum2].coord) <= max_distance:
                            close_pairs.append((rnum1, rnum2))
        if close_pairs:
            found.append((uniprot_id, name, close_pairs, tm_res_ranges))
        m.delete()
    return found, missing

def alphafold_database_model(session, uniprot_id, alphafold_dir):
    filename = f'AF-{uniprot_id}-F1-model_v4.cif'
    from os.path import join, exists
    path = join(alphafold_dir, filename)
    if not exists(path):
        return None
    from chimerax.mmcif import open_mmcif
    s, msg = open_mmcif(session, path)
    return s[0]

def open_entries(session, entries, alphafold_dir):
    models = []
    for uniprot_id, name, close_pairs, tm_res_ranges in entries:
        m = alphafold_database_model(session, uniprot_id, alphafold_dir)
        models.append(m)
```

```
        # Select transmembrane residues
        rnums = residue_numbers_from_ranges(tm_res_ranges)
        for r in m.residues:
            if r.number in rnums:
                r.atoms.selected = True
    session.models.add(models)

uniprot_xml_path = 'UP000001940_6239.xml'
alphafold_dir = 'alphafold_models'
max_distance = 10

ulist = find_uniprot_transmembrane_cysteines(uniprot_xml_path)
print(f'{len(ulist)} UniProt entries')
ntm = len([uniprot_id for uniprot_id, name, paired_hel_cys, tm_res_ranges in ulist if tm_res_ranges])
ntm4 = len([uniprot_id for uniprot_id, name, paired_hel_cys, tm_res_ranges in ulist if
len(tm_res_ranges)>=4])
print(f'{ntm} entries with annotated transmembrane regions')
print(f'{ntm4} entries with 4 or more transmembrane helices')
ntm4p = len([uniprot_id for uniprot_id, name, paired_hel_cys, tm_res_ranges in ulist
            if len(tm_res_ranges)>=4 and len(paired_hel_cys)>=2])
print(f'{ntm4p} entries with 4 or more transmembrane helices and at least two with CC, CxC or CxxC
cysteine pairs')

uclose, missing = check_for_close_cysteines(session, ulist, alphafold_dir, max_distance)
print(f'{len(uclose)} with paired cysteines in two helices closer than {max_distance}A')

entries = []
for uniprot_id, name, close_pairs, tm_res_ranges in uclose:
    rpairs = ' '.join(f'{r1}:{r2}' for r1,r2 in close_pairs)
```

```
    tmranges = ' '.join(f'{r1}-{r2}' for r1,r2 in tm_res_ranges)
    entries.append(f'{uniprot_id},{name},{rpairs},{tmranges}')

print()
print('# UniProt ID, protein name, residue numbers of close paired cysteines, transmembrane ranges')
print('\n'.join(entries))
print()

if missing:
    me = '\n'.join(f'{uniprot_id},{name}' for uniprot_id, name in missing)
    print(f'No alphafold model for {len(missing)} entries with cysteine pairs:\n{me}')

open_entries(session, uclose, alphafold_dir)
```
